# Superior Colliculus Neuronal Ensemble Activity Signals Optimal Rather Than Subjective Confidence

**DOI:** 10.1101/157123

**Authors:** Brian Odegaard, Piercesare Grimaldi, Seong Hah Cho, Megan A.K. Peters, Hakwan Lau, Michele A. Basso

## Abstract

Recent studies suggest that neurons in sensorimotor circuits involved in perceptual decision-making also play a role in decision confidence. In these studies, confidence is often considered to be an optimal readout of the probability that a decision is correct. However, the information leading to decision accuracy and the report of confidence often co-varied, leaving open the possibility that there are actually two dissociable signal types in the brain: signals that correlate with decision accuracy (optimal confidence) and signals that correlate with subjects’ behavioral reports of confidence (subjective confidence). We recorded neuronal activity from a sensorimotor decision area, the superior colliculus (SC) of monkeys, while they performed two different tasks. In our first task, decision accuracy and confidence co-varied, as in previous studies. In our second task, we implemented a novel motion discrimination task with stimuli that were matched for decision accuracy but produced different levels of confidence as reflected by behavioral reports. We used a multivariate decoder to predict monkeys’ choices from neuronal population activity. As in previous studies on perceptual decision-making mechanisms, we found that neuronal decoding performance increased as decision accuracy increased. However, when decision accuracy was matched, performance of the decoder was similar between high and low subjective confidence conditions. These results show that the SC likely signals optimal decision confidence similar to previously reported cortical mechanisms, but is unlikely to play a critical role in subjective confidence. The results also motivate future investigations to determine where in the brain signals related to subjective confidence reside.

**Significance Statement:** Confidence is thought to reflect the rational or optimal belief concerning one’s choice accuracy. Here, we introduce a novel version of the dot-motion discrimination task with stimulus conditions that produce similar accuracy but different subjective behavioral reports of confidence. We decoded decision performance of this task from neuronal signals in the superior colliculus (SC), a subcortical region involved in decision-making. We found that SC activity signaled a perceptual decision for visual stimuli, with the strength of this activity reflecting decision accuracy, but not the subjective level of confidence as reflected by behavioral reports. These results demonstrate an important role for the SC in perceptual decision-making and challenge current ideas about how to measure subjective confidence in monkeys and humans.

## Introduction

When we view the world, our experience often includes an assessment of how confident we are in our perceptual decisions. For example, when driving on a foggy morning, there are moments when we can readily identify elements in our surroundings, and other moments when we are less sure about what lies ahead. Survival in any dynamic environment depends on being able to accurately assess how reliable our perceptions and decisions are in a given instance. Here we ask, how is this subjective sense of confidence in our perceptual decisions represented in the brain?

Work in awake macaques reveals neuronal correlates of confidence in sensorimotor circuits involved in decision-making and action generation, such as the lateral intraparietal area (LIP) (1) and the supplementary eye fields (SEF) (2). One pioneering study of the neurophysiological underpinnings of confidence employed an “Opt-Out” perceptual decision-making task (1). In this task, monkeys made decisions about the primary direction of motion in random dot displays and reported those decisions by making a saccade to one of two targets located in the visual field that corresponded to the dominant dot motion directions (right or left). On some trials, an Opt-Out option appeared orthogonal to the other targets and was associated with a smaller but guaranteed reward; choosing the Opt-Out option indicates less confidence in the decision (3–6). In this task, neurons recorded from area LIP discharged with the highest rates when monkeys correctly chose targets associated with motion toward the response field (RF) and discharged with the lowest rates for correct, opposite RF choices (1). When monkeys chose to Opt-Out, LIP neurons discharged at intermediate levels; these results were interpreted in support of the idea that LIP neurons encode a signal of decision confidence.

An issue arising from this LIP study and most other previous studies of confidence is that decision accuracy and confidence co-vary. That is, since subjects are usually more confident when they perform better on a given task, purported neuronal correlates of confidence may signal decision accuracy rather than subjective confidence *per se.* To make progress, two contributions may be needed: (1) a distinction between different notions of confidence, and (2) a paradigm that dissociates subjective confidence and accuracy. Here, we address both needs.

First, according to one influential theoretical framework (7), confidence can be defined as the probability that a choice is correct, given some available sensory evidence. This probability can be formalized using a binary decision variable *z* which can take on values of left or right, a function *d* which captures the particular choice that is a function of the visual information, and a variable *m* to reflect the *current* choice of left or right. When a choice is made, it can either be correct (when *z = m* and *d = m*) or incorrect (*z = j* and *d = m*, for all *j ≄ m*) (7). Thus, a definition of confidence as the probability that a choice is *correct* given sensory evidence can be written as:

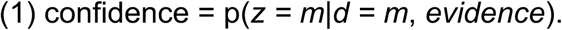

However, as has been noted previously (7), observers’ confidence judgments do not always follow this quantitative definition (8–12). Thus, a distinction must be made between a type of confidence that follows this prediction, and a type of confidence that may deviate from this value empirically.

We introduce the terms “optimal confidence” and “subjective confidence” to refer to these two types of confidence, respectively. “Optimal confidence” refers to the probability in Eq. 1 when it is learned perfectly well: this form of confidence represents the belief of an ideal observer who makes accurate metacognitive judgments about whether a given choice will be rewarded. That is, optimal confidence comes from an ideal observer who accurately computes the probability that a perceptual decision is correct, and therefore will be able to maximize reward if one is to perform the abovementioned Opt-Out task. A neuronal correlate of optimal confidence can be found in population-level activity that effectively distinguishes between conditions yielding different levels of accuracy in a task.

“Subjective confidence” is an actual observer’s internal belief about a perceptual decision, which is potentially prone to error. This form of confidence can deviate from the theoretical ideal of confidence specified by Eq. 1, and can be indexed by behavioral reports which may or may not track decision accuracy perfectly. Because behavioral reports of confidence tend to track optimal confidence at least to some extent, isolating a true neural signal of subjective confidence is difficult. However, this can be facilitated by paradigms that match decision accuracy across conditions, yet yield different behavioral reports about confidence. In this situation, a neuronal correlate of subjective confidence can be found in neural activity that tracks the behavioral reports of varying degrees of confidence, amid constant decision accuracy.

Recent work indicates it is possible to dissociate the capacity to perform perceptual tasks from confidence reports by chemically inactivating the pulvinar (13) or orbitofrontal cortex (14), or psychophysically in humans (15–17). Therefore, we reasoned we could develop visual stimuli that would lead to similar decision accuracy (and therefore, similar levels of “optimal confidence”), but yield different levels of confidence as measured by behavioral reports on individual trials (i.e., “subjective confidence”). Creation of these stimuli would allow us to investigate the neuronal mechanisms of confidence by determining whether activity in a given area signals optimal confidence, subjective confidence, or both (see Fig. S1, which provides a formulation of the two types of confidence in terms of Signal Detection Theory).

Therefore, in this study, monkeys performed two sets of experiments. The first was an Opt-Out task in which decision accuracy co-varied with confidence similar to that performed previously for recordings in LIP (1), allowing us to search for neural correlates of optimal confidence. In the second experiment, building on innovative psychophysical work done in humans (15–18), we introduced a new version of the dot-motion direction discrimination task in which we dissociated reports of confidence from decision accuracy on individual trials. Using this new task, we were able to successfully match decision accuracy (as defined by the signal detection theory measure *d*’ (19– 21)), but produce different levels of confidence (defined as the probability of selecting the Opt-Out target when it was available), so that we could investigate the neuronal correlates of subjective confidence.

As monkeys performed these tasks, we recorded from multiple neurons simultaneously in the superior colliculus (colliculus), a subcortical structure that receives input from LIP and SEF and is involved in decision-making (22–28). We combined these behavioral paradigms and multi-neuron recordings with a machine learning approach (29) to decode population-level activity from hundreds of neurons recorded from the colliculus. We found that in the first task, a population decoder distinguished between high confidence and low confidence trials in much the same way as LIP (1), providing strong evidence that the colliculus contributes to decision-making and optimal confidence in a manner similar to LIP. However, in our novel task in which visual stimuli were matched for sensitivity (*d’*) but resulted in different reports of confidence, population-level activity in the colliculus failed to distinguish between conditions with different degrees of subjective confidence. Together, these findings support the hypothesis that the colliculus signals optimal confidence in dot-motion discrimination tasks, rather than subjective confidence. These results also reveal important considerations for the interpretation of existing data on decision-making confidence in other brain regions, too.

## Results

We used a multivariate decoding approach to assess population-level representations of perceptual decisions and confidence in the superior colliculus using random dot motion discrimination tasks. We had two aims. Our first aim was to determine whether activity measured in the colliculus was similar to that observed previously in area LIP during performance of a confidence task (1). Our second aim was to arbitrate between two competing hypotheses: that neuronal activity in the colliculus primarily signals “optimal confidence,” as signals about confidence may correlate with decision accuracy, or alternatively, that activity in the colliculus signals “subjective confidence,” as neuronal signals may differentiate between conditions where *d’* is matched, but confidence reports vary. We focus here on results obtained from a population decoding method. Further descriptions of neuronal activity and analyses will be reported in detail elsewhere.

We recorded neuronal activity in the colliculus using V-probe laminar electrodes containing 16 recording contacts (see Methods). We measured both single and multi-neuron activity while monkeys performed a dot-motion discrimination task (Fig. 1A, B). Each trial began when the animal established fixation on a central dot. Then, either 2 or 4 choice targets appeared for 500ms. After this delay, the dot motion stimulus appeared at the center of the screen for 200ms. When the motion stimulus disappeared, a delay-period, selected randomly from between 500–600ms, ensued. The fixation dot then disappeared and monkeys indicated their motion direction decision by making a saccade to one of the choice targets, and they received a reward (sip of juice) for correct decisions. Importantly, on some trials there was an Opt-Out option. Choosing this target bypassed the motion discrimination question, and led to a guaranteed but smaller reward compared to that received for correct decisions.

**Fig. 1.**
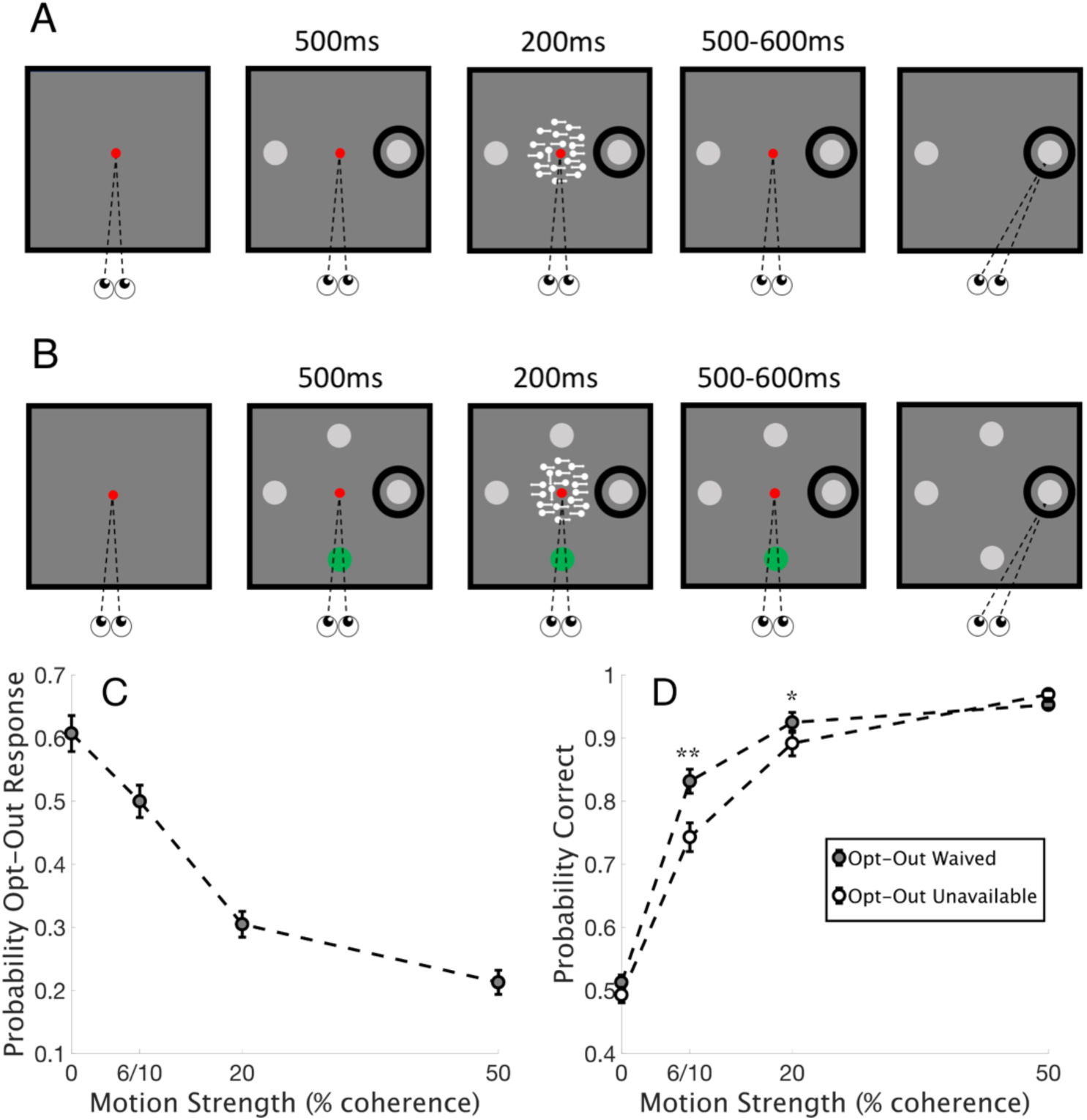
Stimulus-Matched assessment of decision confidence in monkeys. The behavioral task showing a trial in which the Opt-Out option was unavailable (A) and available (B). The trial types shown in A and B were randomly interleaved in each of the 19 Stimulus-Matched sessions. The red dot shows the fixation point, the gray dots show the possible choice targets, and the green dot shows the Opt-Out option. The black circle shows the RF. (C) The probability of choosing the Opt-Out option on trials when it was available (shown in B) is plotted as a function of motion coherence. Circles show means across sessions, and bars show SEM across sessions. Note that monkeys chose the Opt-Out option more often when motion coherence was low, indicating they were less confident about the motion direction decision. The number of trials making up this data set is 14642, as it includes all trials where the Opt-Out was offered. (D) The probability of correct choices is plotted against motion coherence for two monkeys using the same set of data as in C, but now plotting trials where an explicit decision about the motion direction was made (i.e., including trials with the Opt-Out unavailable, and excluding aborted trials, trials where the Opt-Out was selected, and trials where the lateral inhibition target was selected). The number of trials making up this data set is 13346. Circles show means across sessions, and bars show SEM. Gray filled circles show data when the Opt-Out option was available but waived (trials shown in B) and open circles show data when the Opt-Out option was unavailable (trials shown in A). Decision accuracy is higher for intermediate motion strengths when the Opt-Out target was available but waived, presumably reflecting higher confidence (t-tests, Bonferroni corrected, *p < .01, **p < .001).

On trials when the Opt-Out option was available (Fig. 1B), we also included a fourth choice option which was opposite in location to the Opt-Out location to control for possible lateral interactions (see Methods for details). The fourth option never led to reward and was rarely chosen (~6.3% of all trials in Stimulus-Matched sessions). For each session, at least one of the choice targets appeared in the response field (RF) of at least one neuron recorded from the 16 contacts (black circle, Fig. 1A). The two trial types with (Fig. 1B) and without (Fig. 1A) the Opt-Out option available were randomly interleaved; because the properties of the random dot motion stimulus were identical between these trial types, we call these “Stimulus-Matched” sessions.

We reasoned that choices made with the Opt-Out unavailable occur with a mix of high and low confidence, as monkeys are forced to choose one of the two targets. In trials with the Opt-Out option available, monkeys could report their level of confidence: trials in which monkeys chose the Opt-Out target indicate low confidence, whereas trials in which monkeys waive the Opt-Out option and choose one of the targets corresponding to a direction of motion instead, indicate high confidence (1, 3–5).

Figures 1C and 1D show the behavior measured in trials with and without the Opt-Out option available. The probability of selecting the Opt-Out option, when available, decreased as a function of motion coherence, consistent with higher confidence on higher motion coherence trials (all t-tests between conditions p < .05, Bonferroni corrected; Fig 1C). Comparing trials in which the Opt-Out option was available and unavailable shows that at intermediate motion strengths, monkeys have a higher probability of being correct when the Opt-Out option is available and waived compared to when it is unavailable, indicating higher confidence (Fig. 1D).

To determine if neuronal ensemble activity in the colliculus correlates with optimal confidence, we employed multivariate classifiers to evaluate how population-level activity emerged over time as monkeys made decisions in the Opt-Out available and unavailable trials. Previous work shows that LIP discharge rates differ when a correct choice is reported by making a saccade toward the target in the RF (Target-in, or “T_in_”) or away from the RF (Target-out, or “T_out_”) (1). Here, we used a similar approach by evaluating the classifier’s ability to predict correct T_in_ and T_out_ choices with the Opt-Out choice available (but waived) and unavailable.

First, we assessed the area under the ROC curve (AUC) for the classifier (see Methods) to evaluate the degree to which population-level activity in the colliculus may be informative for trial-by-trial predictions of particular behavioral responses (e.g., T_in_ vs. T_out_ choices). This provides us with a measure of how effectively neuronal population activity discriminates between specific perceptual decisions. We focused on a comparison between the Opt-Out Waived condition and the Opt-Out Unavailable condition, because in both conditions the motoric behavior is similar (i.e., the monkeys made saccades to choose one of the options to reflect a perceptual decision rather than the Opt-Out option), and yet optimal confidence is expected to be higher in the Opt-Out Waived condition; higher confidence and expected accuracy presumably cause the monkeys to waive the Opt-Out option when it is available.

Figure 2 shows that neuronal activity in the colliculus signals correct T_in_ and T_out_ choices, and that decoder performance (based on average performance on test sets using 5-fold cross-validation) is higher when the Opt-Out option is available but waived. Here we show the combined results across sessions for two monkeys, but we note that the decoding performance for both monkeys in this task was quite similar (see Figs. S2 and S3). Figure 2A shows that neuronal activity more accurately discriminates correct T_in_ choices vs. correct T_out_ choices when the Opt-Out choice was available but waived, compared to trials where Opt-Out was unavailable (t-tests for all time windows > 230ms after motion onset in middle panel, t(18) > 2.8, p < 0.05). Following previous research (1), we interpret this as evidence that the information contained in the neuronal population activity signals optimal confidence, as the population activity correlates with the differences in decision accuracy across these two conditions. To control for multiple comparisons throughout the entire motion onset period, we used the False Discovery Rate (FDR) method (30) to evaluate significance at each time point. With a false discovery rate of 0.01, while four time windows between 100–230ms were marginally significant, all time windows greater than 230ms after motion onset were highly significant. We also performed further analyses to verify that decoding performance was not just driven by a few single neurons containing strong decision-related activity (as have been identified previously in the colliculus), and that population-level analyses added information over and above what single units can indicate (Fig S4).

We also computed the posterior probabilities for trial-by-trial predictions made by the classifier. Roughly, following previous studies, these probabilities are interpreted as reflecting the strength of the internal decision evidence available to the animal (29–31). We sorted the classifier’s predictions by trial type over time (Figure 2B). Similar to what was shown using the AUC metric, differences between the posterior probabilities between T_in_ and T_out_ correct choices were significantly greater for the Opt-Out Waived trials starting approximately 230ms after motion onset (all time windows > 230ms after motion onset in middle panel, t(18) > 2.8, p < .05).

While the results from decoding T_in_ vs. T_out_ correct trials as a function of Opt-Out availability provide evidence to show that colliculus activity reflects perceptual decisions and their accuracy, does it reflect confidence behavior in any way for it to warrant the label ‘optimal confidence’? As shown in Fig. S5, the absolute magnitude of the strength of this signal decoded from neuronal populations in the colliculus was indeed correlated with actual opt-out rates. However, the relatively small magnitude of this effect suggests that the full story about confidence behavior may be more complicated.

The results described above provide evidence that the neuronal activity in the colliculus contains information about decision-making and decision confidence in much the same way as reported for area LIP (1). However, as noted, the task design used for both the colliculus and the LIP experiments leaves open the possible interpretation that the activity signals decision accuracy (and “optimal confidence”) rather than subjective confidence, since monkeys also performed better on the Opt-Out Waived trials than on the Opt-Out Unavailable trials. Therefore, we created a version of the dot-motion discrimination task in which decision accuracy was matched while confidence varied by manipulating the ratio of “positive evidence” (the amount of motion evidence towards the correct choice) to “negative evidence” (the amount of motion evidence towards the incorrect choice). Previous work shows that while decision accuracy depends upon the ratio of positive to negative evidence, subjective confidence depends upon the overall magnitude of positive evidence (15–18). Thus, we presented monkeys with trials containing *different ratios of positive and negative evidence* to match decision accuracy (defined as perceptual sensitivity, or *d’*, see Methods) across two conditions (Fig. 3A) while attaining different levels of subjective confidence, as measured by their reports of confidence by choosing to opt out or not.

**Fig. 3.**
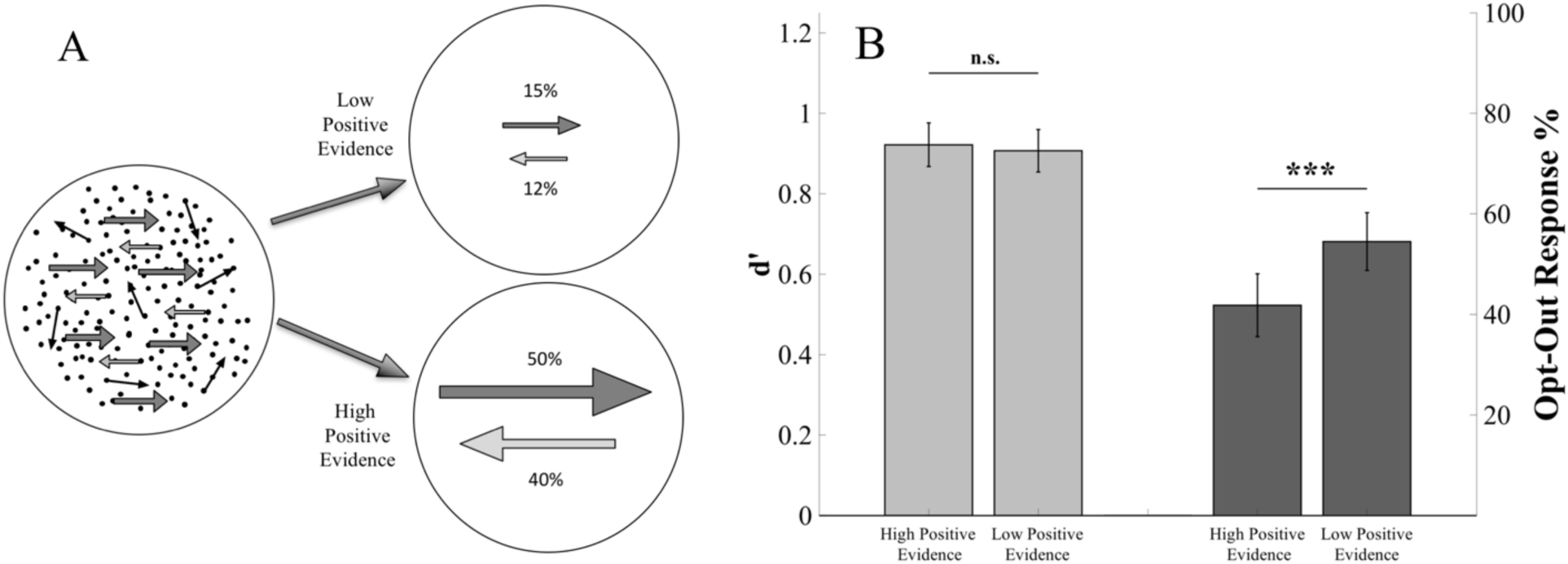
A novel task for dissociating sensitivity and confidence (Sensitivity-Matched) (A) By manipulating the ratio of positive evidence (dot motion toward the correct decision; dark gray rightward arrows) to negative evidence (motion incompatible with the correct decision; light gray leftward arrows), it is possible to match *sensitivity* across two conditions, as measured by *d’*, but achieve different levels of *confidence*, as indexed by the proportion of trials the monkeys chose to Opt-Out. Shown here is a representative example of two conditions that could achieve this result; please note that random dot motion is also included in these conditions, and the exact ratio of positive to negative evidence varied slightly in each session, but the overall number of dots remained constant (see Methods). The sequence of events for this paradigm was identical to the Stimulus-Matched paradigm described in Figure 1, but we refer to this task as “Sensitivity-Matched.” (B) *d’* and the percentage of Opt-Out choices (when the opt-out was available and was selected) are plotted for High Positive Evidence and Low Positive Evidence conditions. Across 23 behavioral sessions from two monkeys, the results show statistically indistinguishable sensitivity (light grey bars) between High and Low Positive Evidence conditions (10197 trials in the *d’* analysis), but different percentages of Opt-Out choices (16471 trials in the opt-out response analysis; dark gray bars, ***p < 10^−5^). Bars show averages across sessions and error bars are SEM.

Figure 3B shows that this manipulation yielded statistically similar levels of decision sensitivity as measured by *d’* (sign test, z = 0.83, p = 0.40), but different degrees of confidence, as indicated by the percentage of trials in which monkeys chose to Opt-Out (sign test, z = −4.59, p < 10^−5^). In this new behavioral task, trials with and without the Opt-Out option were randomly interleaved, allowing us to compute *d’* from trials without the Opt-Out (demonstrating that performance is adequately matched with these stimuli), while evaluating possible differences in subjective confidence from the proportion of trials the Opt-Out was selected when it was available. Data from individual sessions is shown in Figure S6.

With this new task, we evaluated whether the same T_in_ vs. T_out_ decision-related activity differed between the two “Sensitivity-Matched” condition types (High Positive Evidence vs. Low Positive Evidence); that is, whether it passes the test to be considered a neural correlate of *subjective confidence*. Figure 4 shows the decoding results for the Sensitivity-Matched task. The neurons recorded in each of the Sensitivity-Matched sessions used in this decoding analysis were different from the neurons used in decoding the Stimulus-Matched sessions. While trials with and without the Opt-Out were randomly interleaved in the Sensitivity-Matched sessions, we focused our initial decoding analyses solely on the Opt-Out Unavailable trials, excluding Opt-Out Waived trials (Fig. 4). This was to ensure that, should the decoder identify a difference between the two conditions, this difference would not be driven by a sheer difference in internal perceptual response, as the difference in decision criteria for Opt-Out behavior between the High Positive Evidence and Low Positive Evidence conditions means that Opt-Out Waived trials will be more frequent in the High Positive Evidence condition, and as such the average internal response strength will not be matched between the conditions (see Fig. S1).

**Fig. 4.**
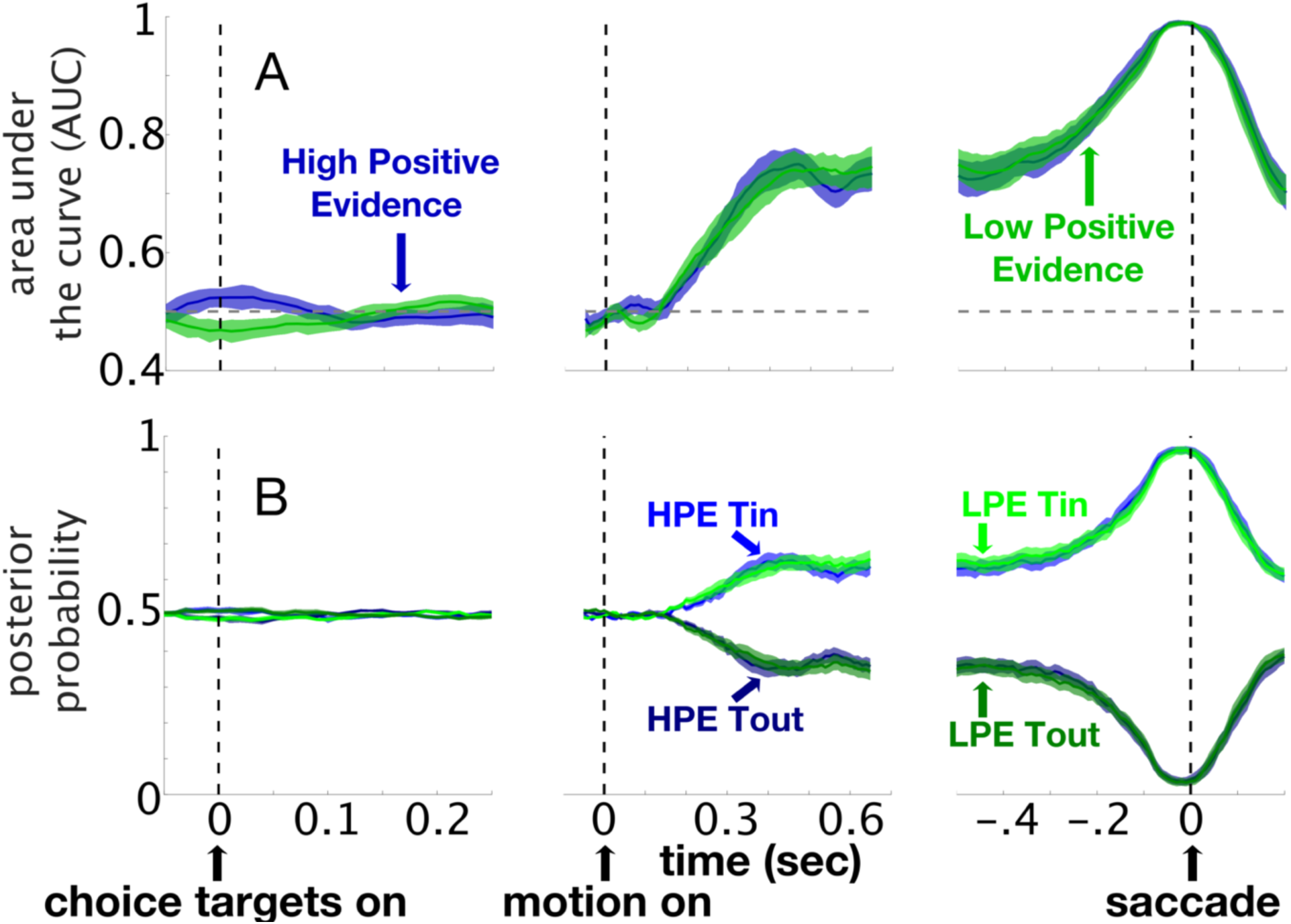
Decoding perceptual decisions made with different levels of confidence and the same level of sensitivity. We trained and tested a decoding model using a 100ms sliding window (step size = 10ms) beginning 50ms before the choice targets appeared through 200ms after the choice report, to predict whether a given correct trial included a saccade toward the choice target in the RF (“T_in_”) or outside of the RF (“T_out_”). The data are from the ‘Sensitivity-Matched’ task shown in Figure 3, and contain 6910 trials from 421 neurons from 2 monkeys (23 total sessions). The decoder was run separately on neurons from each recording session. Each data point represents classification performance of the midpoint of a given 100ms time window (from 50ms before to 50ms after); the figure represents smoothed data using a 5-point moving average. (A) The mean classifier performance as area under the curve (AUC) plotted against time in seconds (sec). (B) The mean posterior probability for T_in_ and T_out_ choices plotted against time in seconds; the y-axis reflects the posterior probability that a given trial contains a correct “T_in_” choice. In all panels, the blue lines and shaded areas show the mean and SEM from the High Positive Evidence (HPE) condition (high confidence) and the green lines and shaded areas show the mean and SEM from the Low Positive Evidence (LPE) condition (low confidence).

Following motion onset, the decoder performance was statistically indistinguishable for both the High Positive Evidence (higher confidence) and Low Positive Evidence (lower confidence) conditions for nearly all time points (59/65 t-tests, t(22) < 2.1, p > 0.05). Importantly, using a false discovery rate of 0.01 to correct for multiple comparisons, none of the time windows reached significance. We observed a similar pattern when comparing the posterior probabilities for the high and low confidence trials from the Sensitivity-Matched task (Fig. 4B). The temporal evolution of the strength of the predictions produced by the classifier were statistically indistinguishable for almost all time points (59/65 t-tests, t(22) < 2.1 p > 0.05), and using the False Discovery Rate method to account for false positives, no significant differences were found for any of the time windows following motion onset.

To further assess whether there are distinct signals for subjective confidence in the activity of colliculus neurons, we performed a cross-generalization analysis. Although the magnitude of decodability between T_in_ and T_out_ choices was similar between the High Positive Evidence and Low Positive Evidence conditions, it is possible that the difference in confidence is reflected by different neurons contributing to this same level of decodability. In that case, some neurons would still meaningfully reflect the different levels of subjective confidence. If that were true, the performance of a classifier that is trained on trials from the High Positive Evidence condition and tested on trials from the Low Positive Evidence condition (or vice versa) should be reduced compared to the performance of classifiers trained and tested within the same condition. That is, if we observe that information was substantially lost through the cross-generalization process, it would provide evidence for distinct neuronal signals for high and low confidence. Figure 5 shows the performance of a classifier trained on trials from the High Positive Evidence condition and tested on trials from the Low Positive Evidence condition as measured by the AUC and posterior probability. This classifier showed similar performance to classifiers trained and tested on trials from a single condition (one-way ANOVA, 64/65 time windows following motion onset, F(66) < 1, p > 0.05); the ability to decode was roughly equivalent across the two conditions, and a comparison between training on Low Positive Evidence and testing on High Positive Evidence yielded statistically indistinguishable results.

Taken together, with differences in population neuronal activity in the colliculus during a Stimulus-Matched confidence task (Fig. 2) but similarity across conditions in a Sensitivity-Matched confidence task (Figs. 4 & 5), these results provide evidence that colliculus activity likely reflects optimal confidence and not subjective confidence *per se*. Although our decoding results suggest that, at the population level, collicular activity discriminates different perceptual decisions at similar levels for the high and low confidence conditions when sensitivity is matched, there may still be individual neurons that reflect subjective confidence in other ways. To investigate this possibility, we computed a normalized ‘discriminability index’ (see Methods) to determine how effectively individual neurons could discriminate between T_in_ and T_out_ choices as a function of confidence in the Sensitivity-Matched task, which included conditions that varied in terms of positive evidence level (High vs. Low), as well as conditions that varied in terms of Opt-Out availability, like in the original task (available vs. unavailable). For a neuron to signal subjective confidence, it should show greater discriminability of T_in_ vs. T_out_ choices not only on trials in which the Opt-Out choice is available but waived (compared to when it is unavailable), but also on trials from the High Positive Evidence condition; put simply, neurons that care about optimal confidence as defined by Opt-Out availability should also care about subjective confidence based on evidence ratios if activity signals subjective confidence at all. Including both of these condition types in the Sensitivity-Matched task allows us to assess this.

**Fig. 2.**
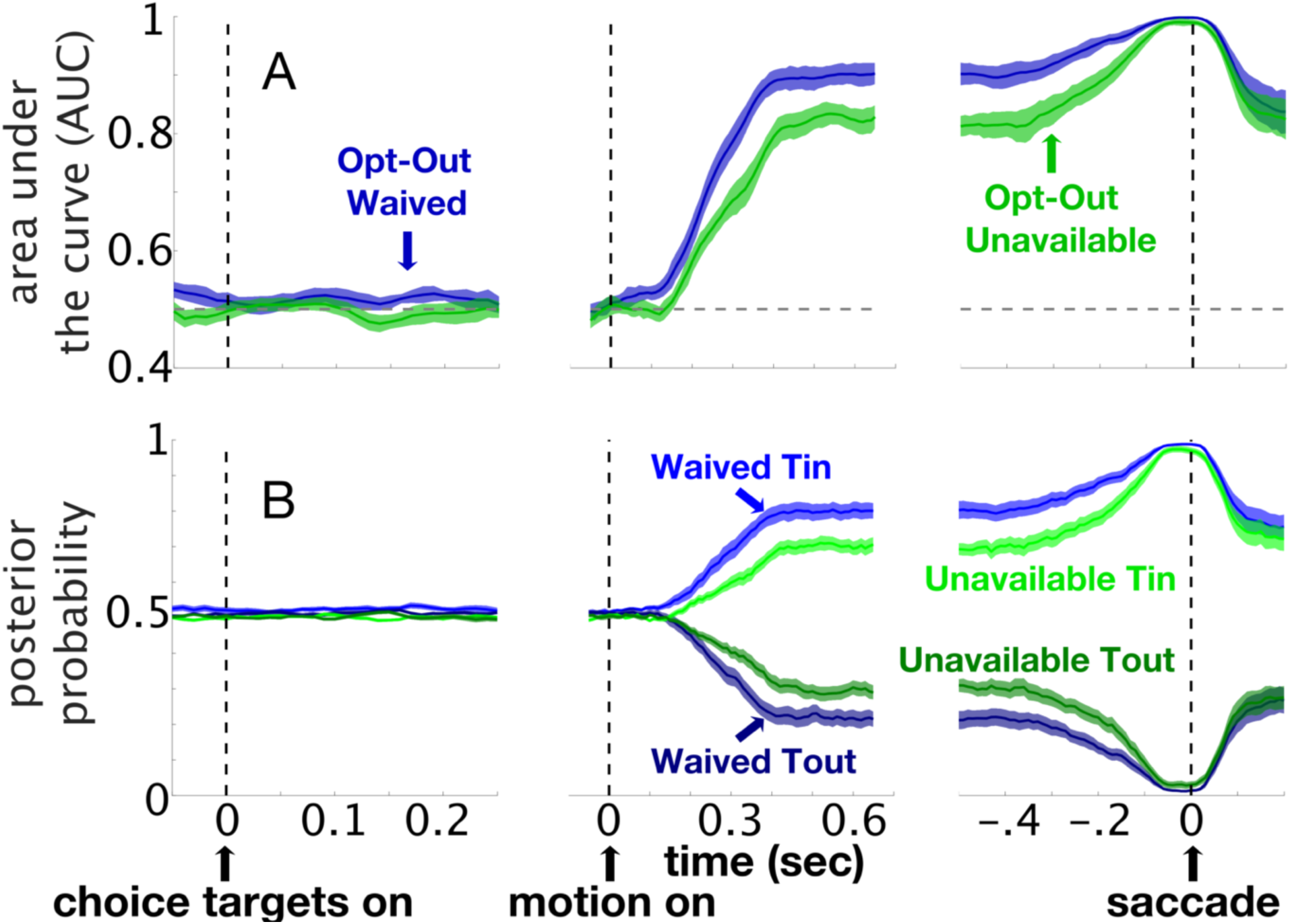
Decoding perceptual decisions made with different levels of confidence for the same motion stimuli (Stimulus-Matched) We trained and tested a decoding model using a 100ms sliding window (step size = 10ms) beginning 50ms before the choice targets appeared through 200ms after the choice report, to predict whether a given correct trial involved a choice in the RF (“T_in_”) or outside of the RF (“T_out_”). 354 collicular neurons were used in this analysis, but the decoder was run independently using 5-fold cross validation on data from each session (which included 9–26 simultaneously recorded neurons, see Methods). The leftmost panels are aligned to the onset of the choice targets, indicated by the dashed vertical line and upward arrow. The middle panels are aligned to the onset of the motion stimulus and the rightmost panels are aligned to the onset of the saccade. Each data point represents classification performance of the midpoint of a given 100ms time window (from 50ms before to 50ms after); the figure represents smoothed data using a 5-point moving average. (A) Mean (thin solid lines) and SEM (shaded areas) classifier performance across sessions shown as the AUC plotted against time for Opt-Out Waived and Opt-Out Unavailable conditions. The ability of the classifier to predict a correct T_in_ or T_out_ choice was better on trials in which the Opt-Out option was available but waived (blue) and monkeys were more confident, compared to when the Opt-Out option was unavailable (green) and monkeys had a mix of higher and lower confidence in their decisions. (B) Similar to A, but plotting the average posterior probability over time. The y-axis is the posterior probability of predicting that a given trial contains a correct “T_in_” choice. This analysis is similar to the “decision variable” used in a previous study (29), and provides an estimate of the strength of the classifier’s predictions.

The discriminability index ranges from −1 to +1. In the Opt-Out available & unavailable conditions (the “Stimulus-Matched” conditions), neurons that maximally discriminate T_in_ and T_out_ when the Opt-Out is available but waived have a value of +1; neurons that maximally discriminate T_in_ and T_out_ when the Opt-Out is unavailable have a value of −1. In the Sensitivity-Matched conditions, neurons that maximally discriminate T_in_ and T_out_ in the High Positive Evidence condition have a value of +1, whereas neurons that maximally discriminate T_in_ and T_out_ in the Low Positive Evidence condition have a value of −1. Neurons with values near 1 on both discriminability indices are neurons that signal confidence.

In line with the decoding results obtained for the original “Stimulus-Matched” task which only included two conditions that varied in terms of Opt-Out availability, we found a significant number of neurons with higher discriminability indices when monkeys waived the Opt-Out choice versus when the Opt-Out choice was unavailable (sign rank test, z = 13.87, p < 10^−42^). For the Sensitivity-Matched conditions, however, discriminability indices regarding capacities as a function of evidence level were distributed symmetrically around 0 (sign rank test, z = −0.78, p = 0.44), indicating a lack of a significant number of neurons that discriminated choices more effectively in the High Positive Evidence trials. Figure S7A shows histograms of the discriminability indices for neurons recorded in Sensitivity-Matched sessions.

Because there were some collicular neurons that gave the appearance of signaling confidence based on their discriminability index (Fig. S7A), we also analyzed neurons with discriminability indices greater than 0 on both measures, to determine if this ability to discriminate T_in_ and T_out_ choices was stable across different trials. We divided each session’s dataset into “odd” and “even” trials, and computed two discriminability indices; one for odd trials and one for even trials for each neuron. If a neuron signals subjective confidence, it should do so for all trials and should do so consistently. If, however, the signal is inconsistent across trials, it is unlikely to provide a signal that is useable by the brain; the odd-even index assesses this. Figure S8A shows the possible confidence neurons-those falling in the upper-right quadrant when the discriminability index for the Stimulus-Matched and Sensitivity-Matched conditions were computed from odd trials only. Computing these same two discriminability indices for the same neurons from even trials (Figure S8B,C), it became clear that while neurons were stable in their increased capacity to discriminate T_in_ and T_out_ as a function of Opt-Out availability (sign rank test, z = 9.82, p < 10^−23^), they were *not* stable in their capacity to discriminate these trial types in Sensitivity-Matched conditions (sign rank test, z = 0.29, p = 0.76). Thus, consistent with the population-level analysis, collicular activity appears better explained by optimal confidence than subjective confidence.

We also assessed whether subjective confidence might be encoded in the colliculus in other ways, beyond what is reflected by the ability to distinguish between correct T_in_ and T_out_ choices. To that end, we analyzed whether we could effectively decode the probability of opting-out in the High Positive Evidence and Low Positive Evidence conditions in our Sensitivity-Matched task. Specifically, in decoding trials when the monkeys chose one of the rewarded targets (both correct & incorrect trials) compared to trials when the monkeys made the decision to opt-out, we could evaluate whether this comparison could effectively discriminate between the High Positive Evidence and Low Positive Evidence conditions, which produced different levels of subjective confidence. Since the purpose here is to actually predict opt-out rate, in this analysis we only included trials where the Opt-Out option was available. Importantly, to make sure the neuronal activity reflected the decision to opt-out rather than just the motoric signal for saccade towards the Opt-Out option, the neurons included here never had the Opt-Out option presented in their response fields. As shown in Fig. S9, this analysis showed that our decoder performed similarly in both the High Positive Evidence and Low Positive Evidence conditions, as both the AUC (64/65 time windows following motion onset, t(22) < 2.1, p > 0.05) and posterior probability (all time windows following motion onset, t(22) < 2.1, p > 0.05) metrics were quite similar across time.

Finally, to further investigate other ways subjective confidence may be coded in the colliculus, we also trained the decoder to directly classify any differences between the High Positive Evidence and Low Positive Evidence conditions (Fig. S10). Despite the various differences in sheer physical stimulus properties and levels of reward expectation, we found that colliculus activity did not strongly distinguish between them.

## Discussion

We combined psychophysics with multi-neuron recordings and population decoding methods to determine whether activity in the superior colliculus of monkeys signals decision confidence. Using a task similar to that used previously in conjunction with recordings in area LIP (1), we identified population-level activity in the colliculus that distinguished between different choices and different levels of confidence in much the same way as LIP. That is, when decision accuracy and decision confidence co-vary, the colliculus signals confidence in a manner similar to LIP. This is consistent with an interpretation that the colliculus, like LIP, signals more than just eye movements, and plays an important role in perceptual decision-making (22, 23, 26, 35–37). However, when comparing collicular activity using a novel task that dissociates optimal from subjective confidence, we found that both population and single neuron activity was indistinguishable between high versus low confidence conditions. Further analyses also failed to find strong evidence in favor of the claim that the colliculus signals subjective confidence *per se*. Thus, we conclude that the role of the colliculus in decision confidence likely primarily concerns optimal confidence.

These findings raise interesting questions regarding previous interpretations of studies. Using an Opt-Out task (1), neuronal correlates of confidence have been found in LIP (1) and in the pulvinar (13). Even though both our study also used the ‘Opt-Out’ design, in a previous investigation (1), monkeys were informed about the Opt-Out option only after the motion stimulus appeared and presumably after they made their decision. In our paradigm, the choice options appeared before the onset of the motion stimulus to avoid visual contamination of the neuronal activity during the stimulus period. Despite this difference, the ability of collicular neurons to distinguish T_in_ and T_out_ choices with different levels of confidence was surprisingly similar to the activity patterns seen in LIP. Similar findings were also obtained in the SEF using a wagering task in which monkeys report their confidence by making ‘bets’ after each perceptual decision (2). To the extent that the colliculus may signal primarily optimal confidence rather than subjective confidence, this open question may apply to those other regions too. Further research is needed to answer this question.

Despite similarities to previous studies, the neuronal ensemble activity in the colliculus did not pass our ‘Sensitivity-Matched’ tests for subjective confidence. However, there could be other neuronal signatures that differ between our two confidence conditions (such as those involving temporal patterns) that our analyses were unable to identify. But to the extent confidence is reflected by firing rate differences between T_in_ and T_out_, as has been assessed by previous studies (1), such activity patterns across the population of neurons assessed seem highly similar between the High Positive Evidence and Low Positive Evidence conditions, as the decoders generalize remarkably well between them (Fig. 5). To exercise further caution, we also conducted analysis of individual neurons (Figs. S7, S8). We found that to the extent that some neurons might show any difference in discriminability between these Sensitivity-Matched conditions, such differences are unlikely to be stable properties of the neurons.

**Fig. 5.**
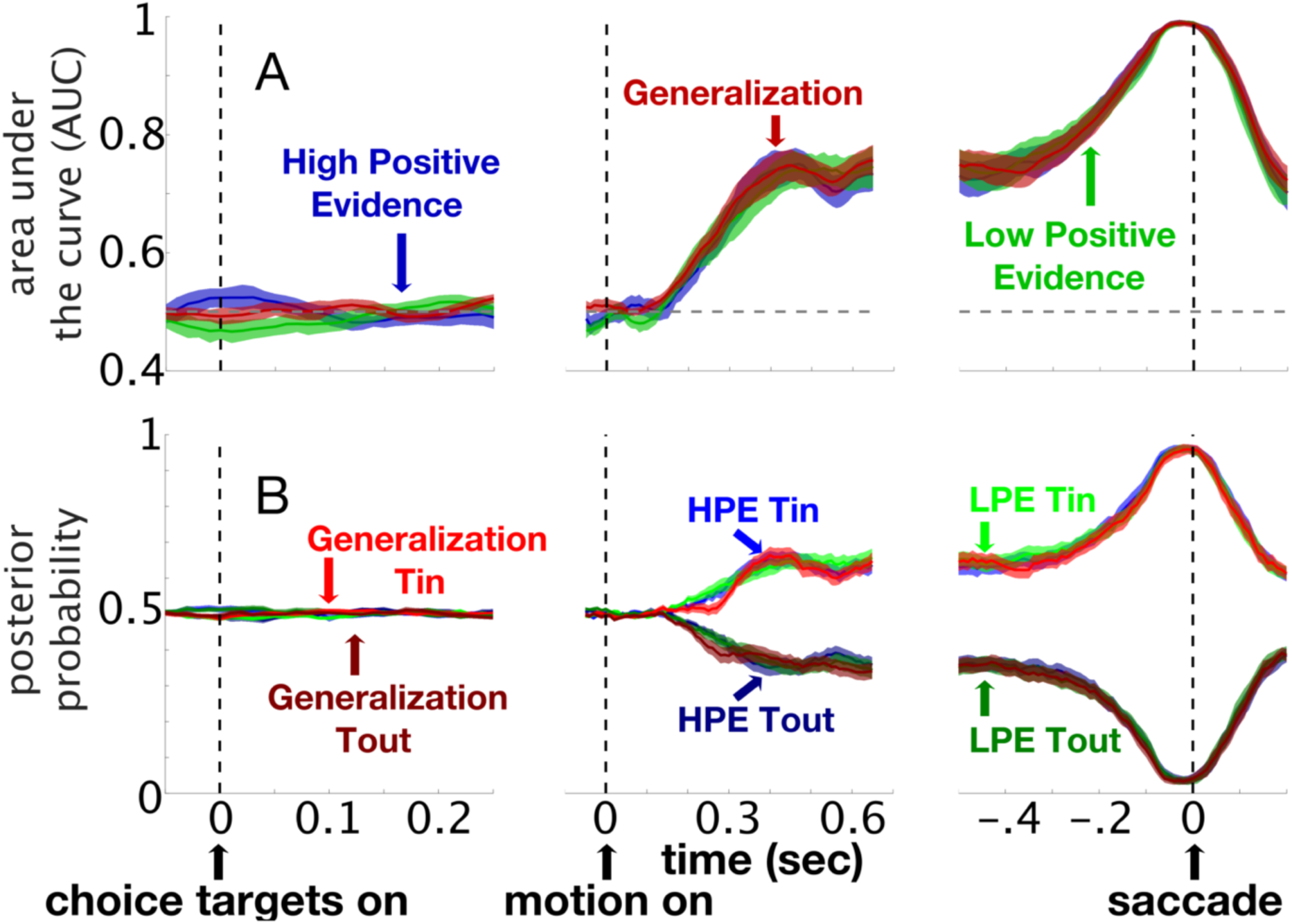
Generalization analysis reveals little evidence for subjective confidence signals in the superior colliculus. Same as in Figure 4, with the addition of results from a linear classifier trained on trials from the High Positive Evidence condition (HPE; high confidence) and tested on trials from the Low Positive Evidence condition (LPE; low confidence), shown in red. Lines show averages and shaded areas show SEM. ‘T_in_’ indicates correct trials in which the monkeys made a saccade toward the choice targets in the RF and ‘T_out_’ indicates correct trials in which the monkeys made saccades toward choice targets outside of the RF.

Based on the human literature (37–39) as well as animal studies (14, 40), one intriguing possibility is that subjective confidence may reside in prefrontal cortex, even under Sensitivity-Matched conditions. Although one previous study (2) recorded from the lateral prefrontal cortex as well as the frontal eye fields, and did not find neurons reflecting *optimal* confidence in these areas as defined above, it remains to be tested whether such neuronal signatures for *subjective* confidence may emerge when confidence is dissociated from sensitivity, or when an ‘Opt-Out’ task rather than a wagering task is adopted. In humans, under Sensitivity-Matched conditions, hemodynamic activity differs between conditions involving different levels of reported confidence (37–39). Applying magnetic stimulation or chemical inactivation to the prefrontal cortex alters confidence reports while sensitivity remains unchanged (14, 38, 40). In another study in monkeys, muscimol injection to the pulvinar impaired confidence reports, as assessed by an ‘Opt-Out’ task, while leaving decision accuracy unchanged (13). Such effects may involve the interactions between the known projections from the dorsal central pulvinar to the prefrontal cortex (41–43). The work in prefrontal cortex and pulvinar, like our work reported here, also argues strongly for a distinction between optimal confidence based on perceptual decisions and subjective confidence that is dissociable from perceptual decisions. We propose that combining our new behavioral task with multi-neuron recordings in the prefrontal cortex and pulvinar may uncover representations of subjective confidence independent of optimal confidence.

Finally, one issue with the Stimulus-Matched task (and Sensitivity-Matched task) is that the condition with mixed high and low confidence trials contains only two visual stimuli, whereas the condition with the opt-out choice available contains four stimuli. It is well-documented that neuronal activity in the colliculus is modulated by choice target uncertainty; specifically, as the number of possible targets increases, activity in the colliculus decreases (22). Our results cannot be explained by this because as the leftmost panels in Figure 2 show, in the opt-out unavailable condition, there are fewer possible stimuli, yet that activity is *indistinguishable* from that seen in the opt-out waived condition in which there are more stimuli on the screen. Moreover, in the middle panel of Figure 2, the activity is *higher* for the opt-out waived condition when there are *more stimuli* on the screen compared to the opt-out unavailable condition when there are fewer stimuli on the screen. This is opposite to what would be expected for an interpretation based on lateral inhibition or uncertainty.

In summary, our findings highlight the important roles played by the superior colliculus in decision-making, beyond its well-known role in eye movements (44), and perhaps more importantly, they raise critical questions about the interpretation of previous findings and open up exciting possibilities for future studies of subjective confidence.

## Methods

### Surgical Procedures

Two male rhesus monkeys (9–13 kg) were prepared for electrophysiological recordings and measurements of eye movements. Anesthesia was induced with an intramuscular injection ketamine (5.0 mg/kg) and midazolam (0.2 mg/kg) and atropine (0.04 mg/kg) was provided to limit salivation. Monkeys were then intubated and maintained at a general anesthetic plane with isoflurane. One hour before the procedure animals received buprenorphine (0.01 mg/kg) and the antibiotic Excede (20 mg/kg; 7 day slow release) and then meloxicam (0.3 mg/kg) at the conclusion of the procedure, and meloxicam (0.2 mg/kg) and buprenorphine (0.01mg/kg) for 3 days post-surgically as analgesia. Monkeys were implanted with MRI compatible headposts and one (monkey H) was implanted with eye loops (45) (46) to measure eye position. In the other monkey (monkey P), eye position was measured with an iView camera (Sensomotoric instruments, Boston, MA). Both monkeys received MRI compatible recording chambers placed over the superior colliculus (AP +3, ML 0) and angled posteriorly at 38°. Precise placement of the post and chambers was performed using MRI-guided surgical software (BrainSight, Rogue Research, Montreal, CA). All surgical procedures were performed under general anesthesia using aseptic procedures. All experimental protocols were approved by the UCLA Chancellor’s Animal Research Committee and complied with and generally exceeded standards set by the Public Health Service policy on the humane care and use of laboratory animals.

### Eye Movement Recording Procedures

We used a QNX-based real-time experimental data acquisition system and windows-based visual stimulus generation system (“Rex” and “Vex”), developed and distributed by the Laboratory of Sensorimotor Research, National Eye Institute in Bethesda, MD (47) to create the behavioral paradigm, display the visual stimulus and acquire two channels of eye position data. Voltage signals proportional to horizontal and vertical components of eye position were filtered (8 pole Bessel-3dB, 180 Hz), digitized at 16-bit resolution and sampled at 1kHz (*National Instruments*; Austin, TX; PCI-6036E). The camera acquired eye position signals were filtered digitally using a built-in bilateral filter. We used an automated procedure to define saccadic eye movements using eye velocity (20°/s) and acceleration criteria (5000°/s^2^), respectively. The adequacy of the algorithm was verified and adjusted as necessary on a trial-by-trial basis by the experimenter.

### Electrophysiological Procedures

We recorded multi-neuron activity from the intermediate layers of the superior colliculus using a platinum/iridium V Probe coated with polyimide (Plexon, Dallas TX) with an impedance of 275 (±50) kΩ. The electrode was aimed at the colliculus perpendicular to its surface using guide tubes positioned with a grid system (48) and advanced using an electronic microdrive system controlled by a graphical user interface (Nan Instruments, Israel). Action potential waveforms were bandpass filtered (250 Hz −5 kHz; 4 pole Butterworth), and amplified, using the BlackRock NSP hardware system controlled by the Cerebus software suite (BlackRock Microsystems, Utah). The voltage data were sampled and digitalized at 30 kHz with 16 bit resolution and saved to disk for offline sorting. For isolating neurons on-line, we used time and amplitude windowing criteria (Cerebus, Blackrock Inc., Utah). Waveforms satisfying these criteria generated TTL pulses indicating the time of occurrence of an action potential and were sampled and digitized at 1kHz with 16 bit resolution and saved to disk.

Action potential waveforms were sorted offline using the Plexon Offline Sorter (Offline Sorter, Plexon, Inc.) and classified into single neurons (n = 115) and multi-neuron (n = 660) activity. At the start of each recording session, we aimed to identify a recording site with at least one buildup neuron, in light of their established role in higher-level phenomena such as attention, selection and decision-making (reviewed in (49)). We classified buildup neurons as those neurons having a significantly higher discharge rate during the stimulus period (200–600ms after motion onset) compared to baseline (200–0ms before the stimulus appears). While the recording procedure first focused on identifying buildup neurons before continuing with the experiment, all neurons that were recorded in a session (both buildup and non-buildup) were used in the decoding analysis for a given session.

Response fields (RF) of collicular neurons were mapped online to provide an estimate of the center of the RF to place at least one choice target. We determined the general characteristics of the neuronal activity and an estimate of the center of the preferred RF by requiring monkeys to make saccades to different locations in the visual field. We made a qualitative assessment on-line about the preferred location on the basis of maximal discharge determined audibly. We confirmed the center of the RF by plotting the discharge as a heat map across visual space. Only neurons with RF eccentricities between 7° and 20° were studied to ensure no overlap of the RF with the centrally-placed moving dot stimulus.

The neurons we recorded from were different in each recording session; the neurons from the 19 Stimulus-Matched sessions which were used were different from the neurons from the 23 Sensitivity-Matched sessions.

### Behavioral Task

We used the same behavioral task in both the Stimulus-Matched and Sensitivity-Matched paradigms. Each trial in both paradigms began when monkeys acquired a centrally-located spot and remained fixated for 500ms. Then, the choice targets appeared. One choice target appeared in the center of the RF of at least one of the recorded neurons (T_in_) and the other choice target appeared in the opposite hemifield (T_out_). These positions were randomized on each trial. For both the Stimulus-Matched and the Sensitivity-Matched paradigms, half of the trials had only two choice targets (i.e., “Opt-Out Unavailable”) and half had an Opt-Out choice target available. These trial types were randomized in each session. All targets, including the Opt-Out, were isoluminant. The location of the Opt-Out choice was orthogonal to the two motion choice targets (90°) and on these trials, we also presented a fourth dot, irrelevant to the task, 180° opposite to the Opt-Out target location. This was included to control for possible lateral interactions (22, 50). That is, to ensure any that differences between the Opt-Out Waived and the Opt-Out Unavailable trials were not driven by introducing an additional response target in an orthogonal location, we introduced a fourth dot to make the stimulus symmetrical, so that each possible target in the Opt-Out available condition was surrounded by a isoluminant targets at the same distance and relative locations.

After the choice targets appeared and monkeys maintained fixation on the central spot for ~500ms, the dot motion stimulus appeared centrally for 200ms. Monkeys maintained fixation for another 500–600ms interval (the exact time was randomly selected between those two times from a uniform distribution), and then were cued to report their decision by removal of the fixation point. If the correct choice occurred, monkeys received a juice reward (0.2ml). If the incorrect choice occurred, monkeys received no reward and a time out of 2000ms. On trials in which monkeys selected the Opt-Out choice, they received a smaller but guaranteed reward (80% of the correct choice reward amount).

### Stimuli

For both tasks, the motion stimulus appeared on a CRT display operating at 60Hz. The motion speed was 5°/s, and the same dots were maintained on the screen for the duration of the stimulus (200ms). Some dots moved coherently in a single direction (coherence percentages described below), while the other dots moved with randomly-selected trajectories. The radius of the motion stimulus was 3°, and the size of dots in the display were 0.05°. The dot density in both tasks was 50 dots per degree squared. Each dot moved in the same direction for the duration of a given trial. For all motion stimuli, the total number of dots appearing on the display was kept constant to maintain isoluminance.

For the Stimulus-Matched paradigm, four motion coherence levels were tested for each monkey. For Monkey P, we tested performance with 20%, 10%, 6% and 0% coherence. For monkey H, we tested performance with 50%, 10%, 6%, and 0% coherence. Different coherence levels were used to yield approximately equivalent performance levels across the two monkeys. Dots moving in random directions were also included, and the total number of dots in all displays was the same.

For the Sensitivity-Matched paradigm, the dot coherence ratios characterized by positive evidence (motion favoring the correct choice) and negative evidence (motion favoring the incorrect choice) were customized for each monkey in each session to yield similar *d’* values across two conditions on trials where the Opt-Out was unavailable, but different amounts of selecting the Opt-Out across those two conditions on trials when it was available. For the eight *d’-matched* sessions for Monkey P, one *d’*-matched session included a 50%PE / 30%NE coherence ratio for High Positive Evidence and 20%PE / 17%NE coherence ratio for Low Positive Evidence; two *d’*-matched sessions included 50%PE / 30%NE coherence ratio for High Positive Evidence and 35%PE / 21%NE coherence ratio for Low Positive Evidence; four *d’*-matched sessions included a 50%PE / 30%NE coherence ratio for High Positive Evidence and 20%PE / 12%NE coherence ratio for Low Positive Evidence; one *d’*-matched session included a 50%PE / 34%NE coherence ratio for High Positive Evidence and 20%PE / 9%NE coherence ratio for Low Positive Evidence.

For the fifteen *d’*-matched sessions for Monkey H, one *d’*-matched session included a 50%PE / 30%NE coherence ratio for High Positive Evidence and 35%PE / 21%NE coherence ratio for Low Positive Evidence; two *d’*-matched sessions included 50%PE / 33%NE coherence ratio for High Positive Evidence and 20%PE / 5%NE coherence ratio for Low Positive Evidence; one *d’*-matched session included a 50%PE / 37%NE coherence ratio for High Positive Evidence and 20%PE / 7%NE coherence ratio for Low Positive Evidence; one *d’*-matched session included a 50%PE / 37%NE coherence ratio for High Positive Evidence and 23%PE / 7%NE coherence ratio for Low Positive Evidence; nine *d’*-matched sessions included a 50%PE / 37%NE coherence ratio for High Positive Evidence and 25%PE / 7%NE coherence ratio for Low Positive Evidence; one *d’*-matched session included 35% PE/ 30% NE coherence ratio for High Positive Evidence and 20%PE / 12%NE coherence ratio for Low Positive Evidence.

### Behavioral Data Analysis

We used signal detection theory to quantify the decision sensitivity of the monkeys in our behavioral task. In this task, monkeys were presented with a dot motion stimulus, and had to make a discrimination judgment as to whether the primary motion direction was to the right or left. *d’* is a measure of an observer’s capacity to perform a sensory task: a *d’* score of 0 indicates a complete inability to discriminate left and right motion directions in this task, while *d’* scores above 0 quantify an observer’s sensitivity to make this type of discrimination. As noted by Wickens (21), *d’* in discrimination tasks can be computed by adding the Z-transformed correct-response probabilities for both stimulus types (p.116). Thus, *d’* was calculated as:

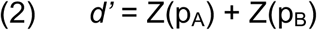

where in this task, p_A_ refers to the probability of a correct judgment for trials where the primary motion direction was towards the left, and p_B_ refers to the probability of a correct judgment where the primary motion direction was to the right. This equation yields the exact same *d’* values as the standard *d’* equation for detection tasks (Z(*Hit Rate*)-Z(*False Alarm Rate*)) but provides a more accurate characterization for discrimination judgments, as “false alarms” are not possible in this type of task, since a primary motion direction is present on every trial.

The Sensitivity-Matched sessions included two different trial types: on some trials, the Opt-Out was unavailable, and monkeys’ only choice was between the two response options. These trials allowed us to determine that our two evidence conditions were matched. On other trials, the Opt-Out was available but could be waived, and these trials allowed us to infer different levels of confidence across these two conditions. We only computed *d’* from trials where the Opt-Out choice was *unavailable*, and focused the decoding analyses on these trials alone. This was done to ensure that, should our subsequent decoding analyses identify a difference across conditions, this difference would not be driven solely by differences in the perceptual criterion used for each condition. The two trial types were randomly interleaved in Sensitivity-Matched sessions, and data from these different trial types is shown in Figure 3 and Figure S4. We also note that in our Sensitivity-Matched sessions, we only analyzed days in which the *d’* scores between our High Positive Evidence and Low Positive Evidence trials were within 0.7 of one another (see figure S4 for individual session results).

### Decoding Analysis

To investigate how population activity in the superior colliculus may be related to optimal and subjective confidence, we applied a decoding model to analyze time-varying neuronal activity, and performed our decoding analyses separately on the data from each recording session. In each session, between 9–26 neurons were recorded from our V Probe recording device, and all units used in decoding for a given session were recorded simultaneously.

We first quantified neuronal discharge rates across all electrodes with a sliding window analysis, computing the sum of action potentials occurring within 100ms time windows (step size=10ms). Next, we applied a logistic regression model using the *fitclinear* function in MATLAB (Mathworks, 2016). The general idea behind this linear classification function is that on any given trial, the overall classification score *f(x)* can be predicted from the neuronal activity at a given time point using the following equation:

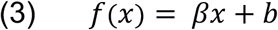

In this equation, *x* is the vector of the summed spike counts for each neuron in a given time window, *β* is a vector representing the linear coefficient estimates for each neuron, and *b* is the scalar bias, reflecting the intercept estimate. However, since our decoding analyses focused on *categorical* outcomes instead of continuous measures, we applied the “logistic” learner from *fitclinear*, which implements the ‘logit’ score transformation function to the raw classification scores to yield the probability of a given class (e.g., *X*), via the following equation:

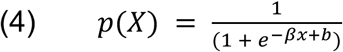

with the following loss function for classification, where *y* ∈{±*1*}:

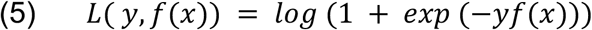

This implementation uses the following ridge regularization penalty to avoid overfitting in our procedure, with a lambda value of (1/number of neurons) in a given session:

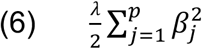

We also implemented a uniform prior in the *fitclinear* function over the two classes, which specified that the two classes being predicted were equally likely on each trial. Finally, we estimated the posterior probabilities for class predictions on each trial using the *predict* function in MATLAB.

As has been noted previously, logistic regression classifiers find the best hyperplane that separates the population response patterns associated with the two classes that are being predicted (29). Therefore, the essential idea behind the aforementioned analysis is that for every trial, the decoder will produce not only a final prediction of, for example, whether the monkey chose the target located in the RF or the target out of the RF, but also a measure of the strength of the prediction (via the posterior probability metric), which corresponds to the prediction’s distance from the hyperplane. We further explain the utility of these metrics in our Results section.

The model was implemented using 5-fold cross validation at each time point, with 80% of the data as the “training set” for fitting the ⍰ coefficients and 20% of the data as the “test set”. In all figures and results, we report the average performance across all five test sets. Two metrics enabled us to assess performance of the model: first, we used area under the ROC curve (AUC) as our method to assess decoder accuracy. Second, we sorted each model’s predictions by trial type, and evaluated the posterior probability of particular class predictions over time. This allowed us to assess the strength of the classifiers’ prediction for each trial type across time, within a range of 0 to 1. Thus, the results we report are based on average AUC and posterior probabilities across the five test sets at each time point.

Three time periods were of particular interest for our decoding procedure. First, the time period around the onset of the targets, to determine whether the pre-stimulus activity held any predictive power for the monkeys’ upcoming decisions. Second, the time period following onset of the motion stimulus, since this is the time when monkeys are forming their decisions and as such, the activity could signal both decision sensitivity and/or subjective confidence. Finally, the time period around the saccade is also informative, as this time window reflects the ceiling for classification performance based on the recorded neuronal activity.

We note that recent work has demonstrated the utility of decoding approaches compared to single-neuron analyses (29, 51), and indeed, our own analysis revealed a stronger capacity to classify correct perceptual decisions by using population-level analyses compared to single neurons (Fig. S3). While we do think that single neuron analysis of our data can also be informative, we think a machine learning approach is particularly advantageous, as decision confidence may be encoded by complex patterns of neuronal activity distributed across many neurons within a brain region, as has been shown in other recent work (39).

### Discriminability Index

In order to assess each neuron’s discriminative capacity for T_in_ and T_out_ choices, we computed a “discriminability index.” This metric produces a normalized value between −1 and 1 specifying both the strength and direction of a neuron’s predictive power for a given two-class discrimination problem. For example, in our initial analysis (see Fig. 2), we classified whether a given correct choice would be toward the RF (T_in_) or away from the RF (T_out_). We hypothesized that the ability to discriminate would change as a function of Opt-Out availability. Thus, we computed the discriminability index for each neuron for the T_in_ vs. T_out_ classification procedures in the following manner:

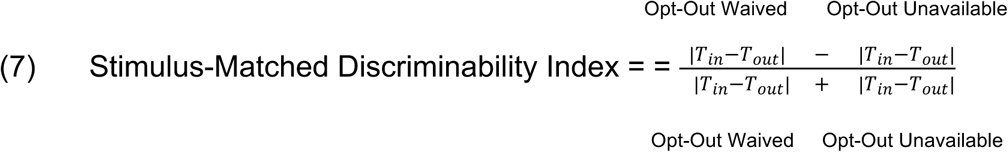

Negative values mean that the neuronal activity is more discriminable for T_in_ compared to T_out_ when the Opt-Out is *unavailable* compared to when it is *waived*; positive values indicate that the neuronal activity is more discriminable between T_in_ compared to T_out_ when the Opt-Out is *waived* compared to when it is *unavailable*. With the sensitivity– matched data, we computed the same discriminability index for trials from the High Positive Evidence condition and trials from the Low Positive Evidence condition using the following equation:

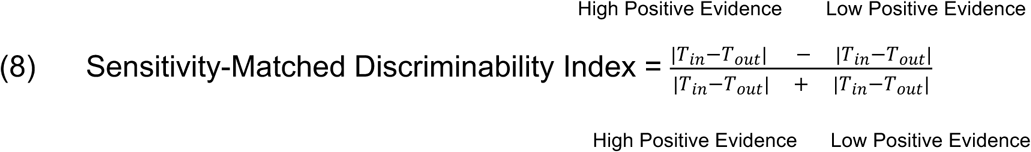

Positive values indicate the neuronal activity is more discriminable for T_in_ compared to T_out_ for trials in the High Positive Evidence condition compared to trials from the Low Positive Evidence condition, and negative values indicate that the neuronal activity is more discriminable for T_in_ compared to T_out_ for trials in the Low Positive Evidence condition compared to trials from the High Positive Evidence condition. We declared neurons to exhibit confidence signals if they fell above 0 on both of these discriminability index metrics (see Results and Fig. S5, S6 for details).

In each experimental session, we computed the discriminability index in each time window from 190ms–650ms after motion onset, and averaged over the discriminability index values to yield a single number for each neuron. This method allowed us to quantify each neuron’s ability to discriminate between the two classes during the main period of evidence accumulation during the trial.

## Acknowledgements

This work was supported by NIH, NS088628 to HL and EY013962 to MAB.

